# Modulation of viral replication, autophagy and apoptosis by induction and mutual regulation of transcription factors EB and E3 during coronavirus infection

**DOI:** 10.1101/2025.04.21.648309

**Authors:** Bei Yang, To Sing Fung, Lixia Yuan, Rui Ai Chen, Ding Xiang Liu

## Abstract

Viral invasion of and replication in cells pose a significant impact on the structure and function of lysosomes. By sensing changes in the lysosome status, cascades of cellular responses are triggered to maintain the lysosomal homeostasis. Two key regulators, transcription factors EB (TFEB) and E3 (TFE3), play essential regulatory roles in these processes by shuttling between the cytoplasm and the nucleus. In this study, we report that infection of cells and/or chickens by gammacoronavirus infectious bronchitis virus (IBV), human betacoronavirus OC43 (HCoV-OC43), and alphacoronavirus porcine epidemic diarrhea virus (PEDV) upregulates the expression of TFEB/TFE3 as well as their downstream targets, and induces the lysosomal stress response. Knockdown of TFE3 alone or together with TFEB demonstrated a pronounced role played by TFE3 in regulating viral replication, virus-induced autophagy and apoptosis in cells infected with these viruses, and a synergistic effect of TFEB and TFE3 in regulation of these processes in cells infected with IBV and HCoV-OC43. Inhibition of the biosynthetic secretory pathway with brefeldin A (BFA) demonstrated that the release of HCoV-OC43 is mainly via the lysosomal pathway. This study provides novel insights into the functional roles of the lysosomal biogenesis and stress response in coronavirus replication and virus-host interactions.

**Importance:** In recent years, more and more studies have shown that lysosomes function more than just as a recycling bin, they also play an important role as a hub for energy and signal transduction, autophagy regulation, and many physiological or pathological processes. This study showed that coronavirus infection activates lysosomal stress responses. As two transcription factors that regulate lysosomal biogenesis, TFEB and TFE3 regulate autophagy and apoptosis during the coronavirus replication cycle. In addition, it has been found that HCoV-OC43 releases viral particles through the lysosomal pathway, which may provide a new reference for the treatment of mitigating coronavirus infection.

## Introduction

Coronaviruses are a class of single-stranded, plus-sense and enveloped RNA viruses that mainly cause respiratory and digestive tract diseases, in various hosts including humans, other mammals (cats, pigs, mice and cattle), poultry and birds (Fung and Liu, 2019a). Similar to other eukaryotic viruses, coronavirus entry into and replication in host cells cause a variety of host cell stress responses, playing essential regulatory roles in viral replication, pathogenesis and cellular responses to viral infection. As a main organelle that senses changes in the intra- and extracellular environments, including nutrient levels and pathogen invasion, lysosomes up-regulate the expression of a number of stress-related genes to allow adaptation of cells to the stimuli and restore homeostasis. This process, known as the lysosomal stress response (Saftig and Puertollano, 2021), is mainly mediated by microphthalmia/transcription factor E (MiTF/TFE). Under stress, MiT/TFE members are activated to coordinate the initiation of multiple stress response pathways, promote the clearance of damaged macromolecules and organelles, maintain cell function, and ultimately restore cell homeostasis.

The lysosomal stress response is mediated by MiTF/TFE transcription factors, and the activation mechanisms of transcription factors EB (TFEB) and E3 (TFE3) have been extensively studied (Raben and Puertollano, 2016). Under physiological conditions, mTORC1 phosphorylates TFEB/TFE3 so that it binds to chaperone protein 14-3-3 and lingers in the cytoplasm. TFEB/TFE3 can also be phosphorylated by a variety of other kinases, including extracellular regulated protein kinases (ERK), glycogen synthase kinase 3β (GSK3β), protein kinase B (Akt) and protein kinase C (PKC), in different cellular signaling pathways (Puertollano et al., 2018). Dephosphorylation and nuclear translocation of TFEB/TFE3 can be triggered by the inactivation of mTORC1 and/or activation of calcineurin/protein phosphatase 2A during starvation and cellular stress ((Martina and Puertollano, 2018). Subsequent binding of the nuclear TFEB/TFE3 to the promoter coordinate lysosome expression and regulation (CLEAR) network would transcriptionally activate autophagy and lysosomal target genes, and regulate apoptosis (Raben and Puertollano, 2016). Depending on the chromosomal maintenance 1 (CRM1) exit, nuclear TFEB/TFE3 can also be re-phosphorylated and relocate back to the cytoplasm (Li et al., 2018; Napolitano et al., 2018; Yin et al., 2020). This phosphorylation-dephosphorylation equilibrium, therefore, regulates the activity of TFEB/TFE3 by controlling their subcellular localization. TFEB/TFE3 can also directly bind and stimulate the expression of tumor necrosis factor-α (TNF-α), interleukin-1 β (IL-1β), IL-6 and other pro-inflammatory factors, playing an important role in the innate immune and inflammatory response (Settembre et al., 2011). Although TFEB and TFE3 are highly similar in amino acid sequence and play similar or even overlapping roles in lysosomal biogenesis and stress response, they may also play distinct regulatory roles in different signaling pathways. One example is that TFEB (but not TFE3) can be specifically phosphorylated by GSK3β in the PKC-GSK3β signaling cascade, differentiating the functional role of TFEB and TFE3 in the lysosomal biogenesis in certain circumstances (Li et al., 2016).

Recent studies have shown that the structure and function of lysosomes are closely related to viral replication, playing particularly important roles in viral invasion of cells and particle release during coronavirus infection. A prerequisite for coronavirus entry into cells is to activate the S protein by protease-mediated cleavage into S1 and S2 subunits before it would be able to mediate viral and cellular membrane fusion (Fung and Liu, 2019a). In addition to direct entry into cells after activation by transmembrane protease serines (TMPRSS2) on the plasma membrane, most coronaviruses enter the endosomal pathway via endocytosis followed by activation of S protein by lysosomal cathepsin L/B, mediating membrane fusion (Pišlar et al., 2020). Reduction of the lysosomal acidity by treatment of cells with inhibitors or knockdown of the lysosomal mass pump ATPase H^+^ transporting V0 subunit D1 (ATP6V0D1) has a significant inhibitory effect on coronavirus replication (Gorshkov et al., 2021). Recent studies have also shown that the release of betacoronavirus particles is mainly through the lysosomal pathway, a process markedly different from other enveloped viruses that utilize the normal cellular secretory pathway (Ghosh et al., 2020).

In this study, we report the upregulation of TFEB and TFE3 in cells infected with avian gammacoronavirus infectious bronchitis virus (IBV), human betacoronavirus-OC43 (HCoV-OC43), and alphacoronavirus porcine epidemic diarrhea virus (PEDV), respectively. Characterization of the functional roles of TFEB and TFE3 in cells infected with these viruses suggests that these two key molecules play important regulatory roles in viral replication, autophagy and apoptosis. In addition, the lysosomal pathway is exploited by HCoV-OC43 in the release of mature virions during their replication cycles. This study provides novel insights into the functional roles of the lysosomal stress response and lysosomal biogenesis in coronavirus-host interactions.

## Material and method

### Cell culture, SPF Chickens and virus

HeLa, HEK293T and Vero cells were cultured in Dulbecco’s modified Eagle’s medium (DMEM, Life Technologies, Carlsbad, CA, USA) supplemented with 10% fetal bovine serum (FBS), 100 U/ml penicillin and 100 mg/ml streptomycin (Gibco, Grand Island, NY, USA). H1299 cells were cultured in RPMI 1640 medium (Gibco) supplemented with 8% fetal bovine serum (FBS), and 100 U/ml penicillin and 100 mg/ml streptomycin (Gibco). All cells were grown in a 37°C incubator supplied with 5% CO_2_.

White Leghorn specific-pathogen-free (SPF) chicks (one-day-old) were obtained from the SPF Experimental Animal Center of Xinxing Dahua Agricultural, Poultry and Egg Co., Ltd. (Xinxing, China), with approved number: SCXK (Guangzhou, China) 2018-0019. The SPF chicks were raised in individually ventilated cages within the SPF animal house of Zhaoqing Dahuanong Biology Medicine Co., Ltd. (Zhaoqing, China).

The egg-adapted Beaudette strain of IBV (ATCC VR-22) was obtained from the American Type Culture Collection (ATCC) and adapted to Vero cells as previously described (Lim and Liu, 1998; Shen et al., 2003). This Vero-adapted strain was named IBV-p65, and the complete genome sequence was uploaded (accession No. DQ001339) (Fang et al., 2005). HCoV-OC43 (accession No. KU131570.1) (Morfopoulou et al., 2016) were also obtained from ATCC, and the complete genome sequences were uploaded. PEDV virulent strain DR13 (PEDV-vDR13) was isolated in Korea in 1999 (accession No. JQ023162) as previously reported (Park et al., 2012).

Virus stock was prepared by infecting monolayers of Vero or H1299 cells with at a multiplicity of infection (MOI) of approximately 0.1 and cultured in DMEM at 37°C for 24 h. After three freeze-thaw cycles, total cell lysates were clarified by centrifugation at 1,500g at 4°C for 30 min. The supernatant was aliquoted and stored at −80°C as a virus stock.

The titers of the virus stocks were determined by 50% tissue culture infective dose (TCID50) assay. IBV, PEDV, and HCoV-OC43 stocks were used for infection of cells at an MOI of ∼2 in all experiments, and the mock control was incubated with same amounts of UV-inactivated virus as previously described (Yuan et al., 2021). Virus infection was carried out by incubation of cells with a virus stock for about 2 h, washing twice with PBS to remove the unbound viruses, and further incubation for the indicated times in each experiment after addition of fresh medium.

### Antibodies, chemicals, and reagents

Anti-β-actin (catalog number HC201-01) and anti-DYKDDDDK (catalog number HT201-01) were purchased from TransGen Biotech (Beijing, China). Anti-TFEB (catalog number37785), anti-TFE3 (catalog number14779S) and anti-PARP (catalog number 9532) were purchased from Cell Signaling Technology (CST, Boston, MA,USA). Polyclonal antibodies against IBV N, M and S proteins were prepared from rabbits immunized with bacterially expressed fusion proteins as previously described (Yuan et al., 2021). Anti-OC43 N was purchased from Sinobiological (Beijing, China).

Nuclear and cytoplasmic protein extraction kit (catalog number P0028) was purchased from Beyotime (Shanghai, China). Brefeldin A (catalog number S7046) was purchased from Selleck (Houston, TX,USA).

### Plasmid construction and transfection

Human TFEB complementary DNA (cDNA) (NM_007162) was amplified from total RNA extracted from H1299 cells by reverse transcription-PCR (RT-PCR), using primer pair CGATGATAA GTCCGGATCCGCGTCACGCATAGGGTTG and AAGATCTGGTACCGAGCTCCTGCAGTCAC AGCACATCGCCCTC. Human TFE3 cDNA (NM_006521) was amplified using primer pair CGATG ATAAGTCCGGATCCTCTCATGCGGCCGAACCAG and AAGATCTGGTACCGAGCTCCTGCA GTCAGGACTCCTCTTCCATGCTGAAGC. Chicken TFEB cDNA (NM_001030922.1) was amplified from chicken tissue RNA extracted using primer pair CGATGATAAGTCCGGATCCGCGTCCCGCATCGGGCTG and AAGATCTGGTACCGAGCTCCTGCAGTCACAGCATGTCTGC GTCCTCC. These PCR products were inserted into pXJ40-Flag linearized with *BamH*I and *Pst*I by homologous recombination, and were named pXJ40-Flag-TFEB, pXJ40-Flag-TFE3 and pXJ40-Flag-TFEB(CHK), respectively.

Plasmid DNA was transfected into cells using TransIntro EL transfection regent (Transgen biotech). Briefly, cells seeded on a 12-well plate were cultured overnight in medium containing 10% FBS, and transfection was performed when cells approached 60-70% confluence, with 2μg of plasmid DNA and 4μl of TransIntro EL diluted each with 100μl of Opti-MEM (Gibco). After mixing by brief vortex and further incubation for 15 min, the mixture was added to each cell well dropwise, and was replaced with normal fresh complete medium after incubation for 4-6 h. At 24 h post-transfection, cells were infected with IBV and harvested at indicated time points for further analysis.

### RNA extraction and RT-qPCR analysis

Total RNA was extracted using the TRIzol reagent (Invitrogen, Carlsbad, CA, USA) and reverse-transcribed using the 5 ×FastKing gDNA dispelling RT SuperMix kit (Tiangen, Beijing, China) according to the manufacturers’ instructions. The first-strand cDNA was then diluted 20-fold with RNase-free water for quantitative PCR (qPCR) analysis using Talent qPCR PreMix SYBR green kit (Tiangen) with a QuantStudio 3 real-time PCR system (Applied Biosystems). The standard protocol included enzyme activation at 50°C for 3 min and an initial denaturation step at 95°C for 3 min, followed by 40 cycles of denaturation (95°C for 5 s) and annealing/extension (60°C for 30 s), with fluorescence acquisition at the end of each cycle. The results obtained were in the form of cycle threshold (CT) values. Using the Δ Δ CT method, the relative abundance of a transcript was calculated using glyceraldehyde-3-phosphate dehydrogenase (GAPDH) as internal control for normalization.

The sequences of qPCR primers are human-GAPDH, CTGGGCTACACTGAGCACC and AAG TGGTCGTTGAGGGCAATG; IBV gRNA, GTTCTCGCATAAGGTCGGCTA and GCTCACTAAA CACCACCAGAAC; human-SQSTM1, CAACATGGTGCACCCCAATGT and CGCTACACAAG TCGTAGTCTGG; human-TFEB, CAAGGCCAATGACCTGGAC and AGCTCCCTGGACTTTT GCAG; human-TFE3, GCTGCTTTCCTTGGC and ATCTGAGGGCGGTGC; human-M6PR, CTC AGTGTGGGTTCCATCTTAC and GGGAAACTGCTCCATTCCTT; human-CTSB, GGACAAGC ACTACGGATACAA and GTAGAGCAGGAAGTCCGAATAC; human-RAB7A, CCTGGAGTCTT GGCCATAAAG and GAGAAGGTCCAAGTTCTGGTTC; human-MCOLN1, GGAAAGCAGCT CCAGTTACA and GATGAGGCTCTGGAGGTTAATG; human-CHOP, GGAAACAGAGTGGTC ATTCCC and CTGCTTGAGCCGTTCATTCTC; chicken-CTSB, GCACTACGGCATCACATCCT and AACCTGCTCCCCTGACACAT; chicken-GAPDH, GACCACTGTCCATGCCATCA and TTTCCCCACAGCCTTAGCAG.

### Construction of stable knockdown cell clones and RNA interference

Lentivirus-based shRNA vector pLKO.1 was used to infect H1299 cells and the knockdown cell clones were screened with 2μg/ml of puromycin. The sequences of the shRNA sense strands are ATTG TTGCTGACATAGAATTA for TFE3, ACTACCGTTGTTATAGGTGT for negative control. The siRNA duplexes used for RNA interference were purchased from Sangon Biotech (Shanghai, China). The sequences of the siRNA sense strands are AGACGAAGGUUCAACAUCAdTdT for TFEB and GCUGACCCUGAAGUUCAUCdTdT for EGFP as the negative control. Transfection of siRNA was performed using the TransIntro EL transfection reagent (TransGen Biotech) as previously described(Li et al., 2022).

### SDS-PAGE and Western blot analysis

Cells were collected and centrifuged at 16,000g for 3 min. The supernatants were collected and an appropriate amount of 5× sodium dodecylsulfate (SDS) loading dye (Beyotime) was added, cell pellets were mixed with RIPA buffer (Beyotime) and 5× SDS loading dye, boiled at 95℃ for 5 min. After a short centrifugation, same amounts of protein samples were loaded into each well and separated by SDS-polyacrylamide gel electrophoresis (SDS-PAGE). The resolved proteins were then transferred to a 0.2μm nitrocellulose membrane using the Bio-Rad Trans-Blot protein transfer system, the membrane was incubated with 5% skim milk in Tris-buffered saline–Tween 20 (TBST) buffer (20 mM Tris-HCl [pH 7.4], 150 mM NaCl, 0.1% Tween 20) at room temperature for 1-2 h, washed with 1×TBST three times, and incubated with a specific primary antibody dissolved in TBST with 3% (wt/vol) BSA at 4°C overnight. After removal of the primary antibody and washing with 1×TBST, the membrane was incubated with a 1:10,000-diluted appropriate secondary antibody at room temperature for 2 h. Fluorescence imaging was performed using the Azure c600 imager, and densitometric measurement was performed using AzureSpot software. All experiments were repeated at least three times and one of the representative results is shown.

### Nuclear and cytoplasmic protein extraction

Confluent HeLa cells infected with IBV were washed with PBS at different time points post-infection, and cell pellets were collected. Nuclear and cytoplasmic protein extraction experiments were performed using the nuclear and cytoplasmic protein extraction kit (Beyotime), added to 5×SDS, heated at 95℃ for 3-5 min, and analyzed by SDS-PAGE and Western blot.

### Cell viability assay

Cell proliferation experiments were performed using TransDetect Cell Counting Kit (CCK, TransGen Biotech, Beijing, China). Cells were placed in 96-well plates with 5×10^3^ cells in 100μl per well and incubated overnight, 10μl of CCK Solution were added. After incubation for 2-4 h, the absorbance at 450nm was measured by spectrometry in tetraplicate per cell clone.

### Infection of chickens with IBV

One-day-old SPF chickens were randomly divided into two groups (n=5/group), one group infected with (∼105.5 EID50) KP-GI-19, a local IBV isolate of G1-19 genotype(Xiong et al., 2024), by the nasal–ocular route. The remaining five chickens were infected with the same volume of PBS as control. All chickens were euthanized at 7 days post-inoculation for the pathological autopsy and examination of lesions in all organs. Total RNA extracted from each organ was collected for RT-qPCR, and the expression levels of IBV-gRNA and CTSB in all chicks were detected.

### Statistical analysis

One-way analysis of variance (ANOVA) was used to analyze significant differences between the indicated samples and the respective control samples. Significance levels are presented by the P value (ns, nonsignificant; *, P < 0.05; **, P < 0.01; ***, P < 0.001).

## Results

### IBV infection stimulates the nuclear translocation of TFEB and TFE3

The functional effect of IBV infection on TFEB/TFE3 was first assessed by isolating cytoplasmic and nuclear proteins from IBV-infected HeLa cells at different time points post-infection. Western blot analysis showed that TFE3 protein in the cytoplasm gradually decreased and translocated to the nucleus, with increase of the infection time (Fig. 1A). By constructing and utilizing a lysosomal stress reporting system, a HeLa cell clone stably expressing TFEB-EGFP, the distribution of green fluorescence was observed by fluorescence microscopy at different time points post-infection. Nuclear translocation of the majority of the TFEB green fluorescent protein to the nucleus was observed in the infected cells, compared to cells incubated with the UV-inactivated virus at 24 hpi (Fig. 1B). These results suggest that IBV infection indeed induces the nuclear translocation of TFEB and TFE3.

**Figure 1.**
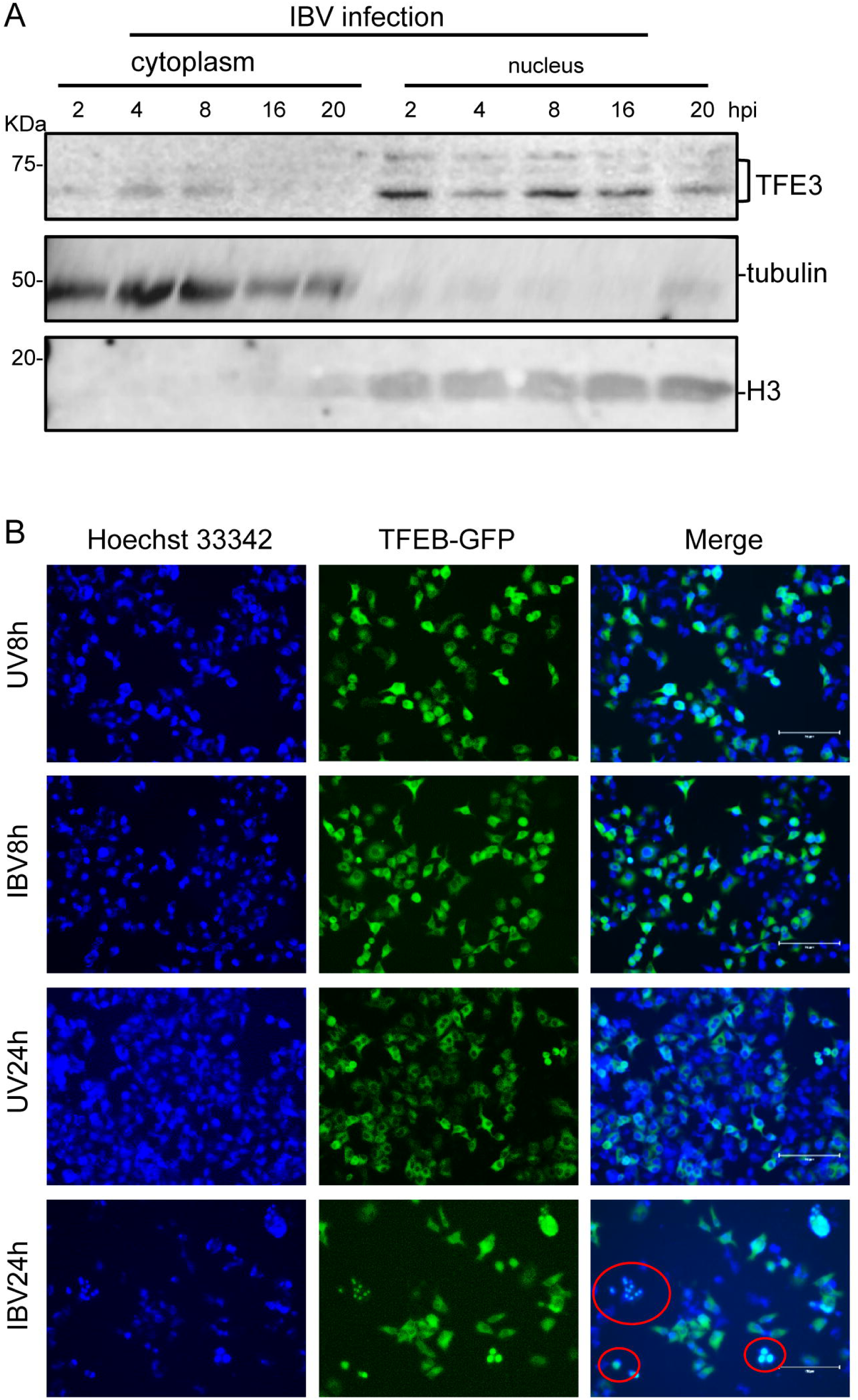
Stimulation of the nuclear translocation of TFEB and TFE3 by IBV infection. A. HeLa cells were infected with IBV at an MOI∼2 and harvested at indicated times post-infection. Nuclear and cytosolic fractions were prepared and subjected to Western blotting with the indicated antibodies. Beta-actin was included as the loading control. B. HeLa cells stably expressing TFEB-EGFP were infected with IBV at an MOI∼2 or mock-treated with UV-inactivated IBV, nuclear stained with Hoechst 33342 at specified time points, and examined by fluorescence microscopy. The merged images show the nuclear colocalization of TFEB. Scale: 75μm.

### The expression of TFEB/TFE3 and target genes is induced by virus infection of culture cells and chicks

To test if the IBV-induced TFEB/TFE3 nuclear translocation activates the lysosomal stress response and up-regulates the expression of the downstream genes, time-course experiments were conducted in H1299 and HeLa cells infected with IBV, and the expression of these genes was analyzed by Western blot and RT-qPCR. Cell starvation can also inactivate mTORC1, resulting in TFEB/TFE3 dephosphorylation from 14-3-3 and nuclear translocation. To exclude this effect, protein and RNA samples were harvested from cells treated with UV-inactivated IBV and infected with IBV, respectively, in the early (8 h post-infection, hpi) and late (20 hpi) stages of the IBV infection cycle. We observed that the protein level of TFE3 in IBV-infected cells was upregulated at both time points, compared with that in cells treated with UV-inactivated IBV, demonstrating that IBV infection of cells indeed upregulated the expression of host TFE3 gene (Fig. 2A).

**Figure 2.**
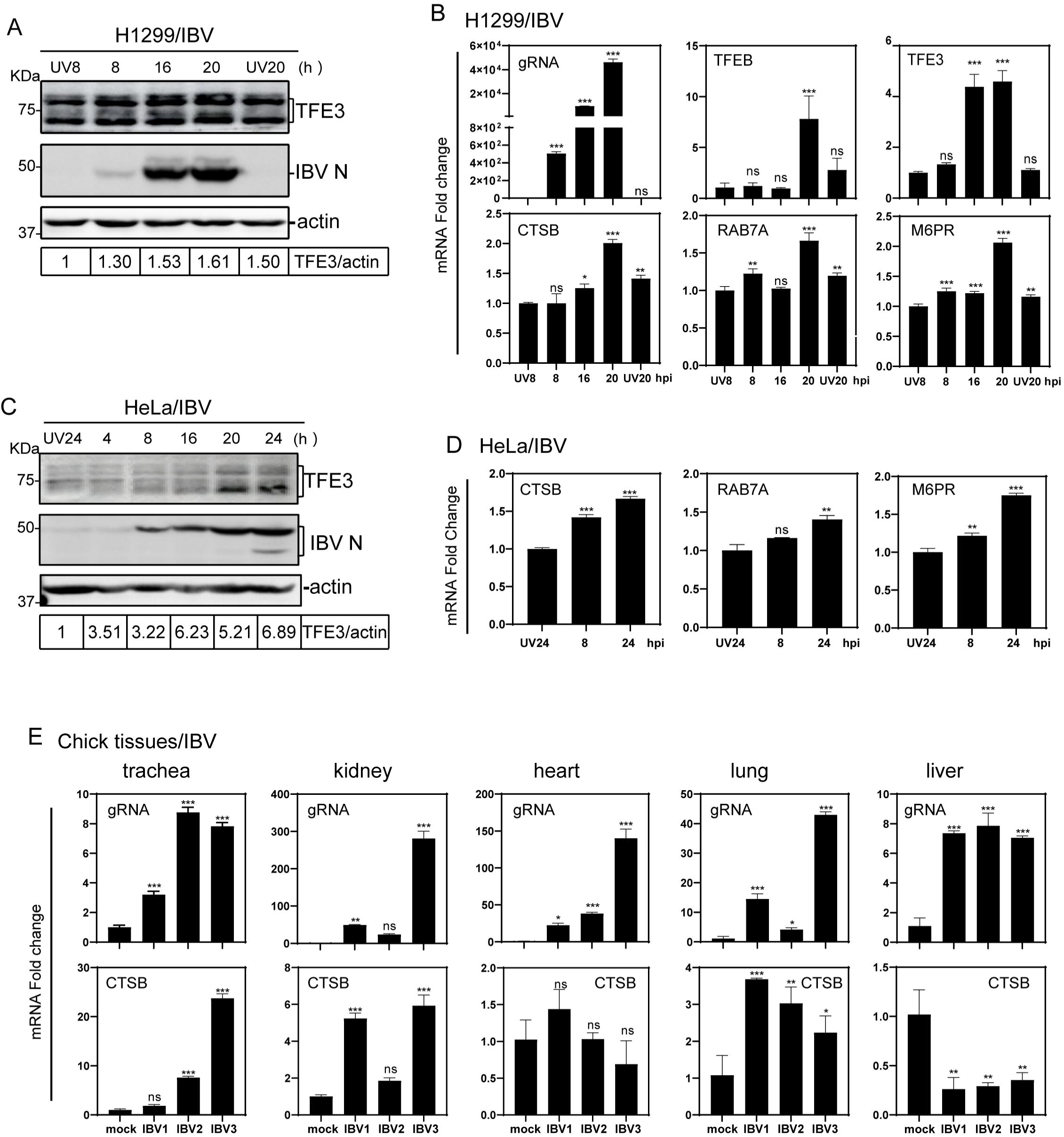
Induction of TFEB/TFE3 and target genes by virus infection of culture cells and chicks. A. H1299 cells were infected with IBV at MOI∼2 or mock-treated with UV-inactivated IBV and harvested at the indicated time points. Total cell lysates were prepared and subjected to Western blot analysis with indicated antibodies. B. H1299 cells were infected and harvested as described in A. Total RNAs were extracted, the level of IBV genomic RNA (gRNA), and mRNA levels of TFEB,TFE3 and downstream target genes CTSB, RAB7A and M6PR were determined by RT-qPCR with the ΔΔCT method after normalization to the GAPDH mRNA in cells treated with UV-inactivated IBV. C. HeLa cells were infected with IBV at MOI∼2 or mock-treated with UV-inactivated IBV and harvested at the indicated time points. Total cell lysates were prepared and subjected to Western blot analysis with indicated antibodies. D. HeLa cells were infected and harvested as described in A. Total RNAs were extracted, the level of IBV genomic RNA (gRNA), and mRNA levels of TFEB, TFE3 and downstream target genes CTSB, RAB7A and M6PR were determined by RT-qPCR with the ΔΔCT method after normalization to the GAPDH mRNA in cells treated with UV-inactivated IBV. E. One-day-old SPF chickens were either infected with a local isolate of IBV strain (KP-G1-19, a G1-19 genotype), or injected with PBS as negative control. At seven days post-infection, trachea, kidney, heart, lung and liver were collected and total RNA extracted. The levels of IBV gRNA and CTSB mRNA in each organ were detected by RT-qPCR with the ΔΔCT method using GADPH from the PBS-injected groups as the standard (ns, non-significant; *, P < 0.05; **, p < 0.01; ***, p < 0.001).

To investigate if IBV-induced nuclear translocation of TFEB/TFE3 activates the CLEAR network, three downstream gene targets, CTSB (cathepsin B) encoding a lysosomal cysteine protease, RAB7A (member of RAS oncogene family) encoding an endolysosomal trafficking small GTPase, and M6PR (mannose-6-phosphate receptor, cation dependent), were analyzed by RT-qPCR. The expression of these three lysosome-related genes was upregulated with increase of the infection time (Fig. 2B). A similar result was also obtained in IBV-infected HeLa cells (Fig. 2C-D).

To reflect the results under physiological conditions in living birds, the expression levels of IBV-gRNA and CTSB in different organs from chickens infected with IBV strain KP-GI-19 were analyzed by RT-qPCR. As shown in Figure 2E, the mRNA level of CTSB gene in the trachea, kidney, heart and lung from infected chicks was significantly up-regulated, compared with that in the control group. Among these organs, the highest is in trachea, with about 25-fold upregulation (Fig. 2E). Taken together, these results demonstrate the induction of lysosomal stress response by IBV infection of culture cells and chickens under experimental conditions.

### Lysosomal stress response is also induced in culture cells infected with two other coronaviruses

To investigate if the lysosomal stress response was also induced in cells infected with other coronaviruses, H1299 cells were infected with coronaviruses HCoV-OC43 and PEDV, respectively, and protein and RNA samples were collected at different time points post-infection for Western blot and RT-qPCR analyses. At the protein level, TFE3 was upregulated along with the increased detection of coronaviral N protein in the time-course experiments (Fig. 3A). At the mRNA level, TFEB, TFE3 and lysosomal target genes MCOLN1, CTSB and RAB7A were also upregulated to varying degrees (Fig. 3B). These results confirm the induction of lysosomal stress response in cells infected with these viruses. The upregulation of these gene expression at varying levels and at different time points of their infection cycles may reflect distinct characteristics of the replication and infection kinetics of individual viruses.

**Figure 3.**
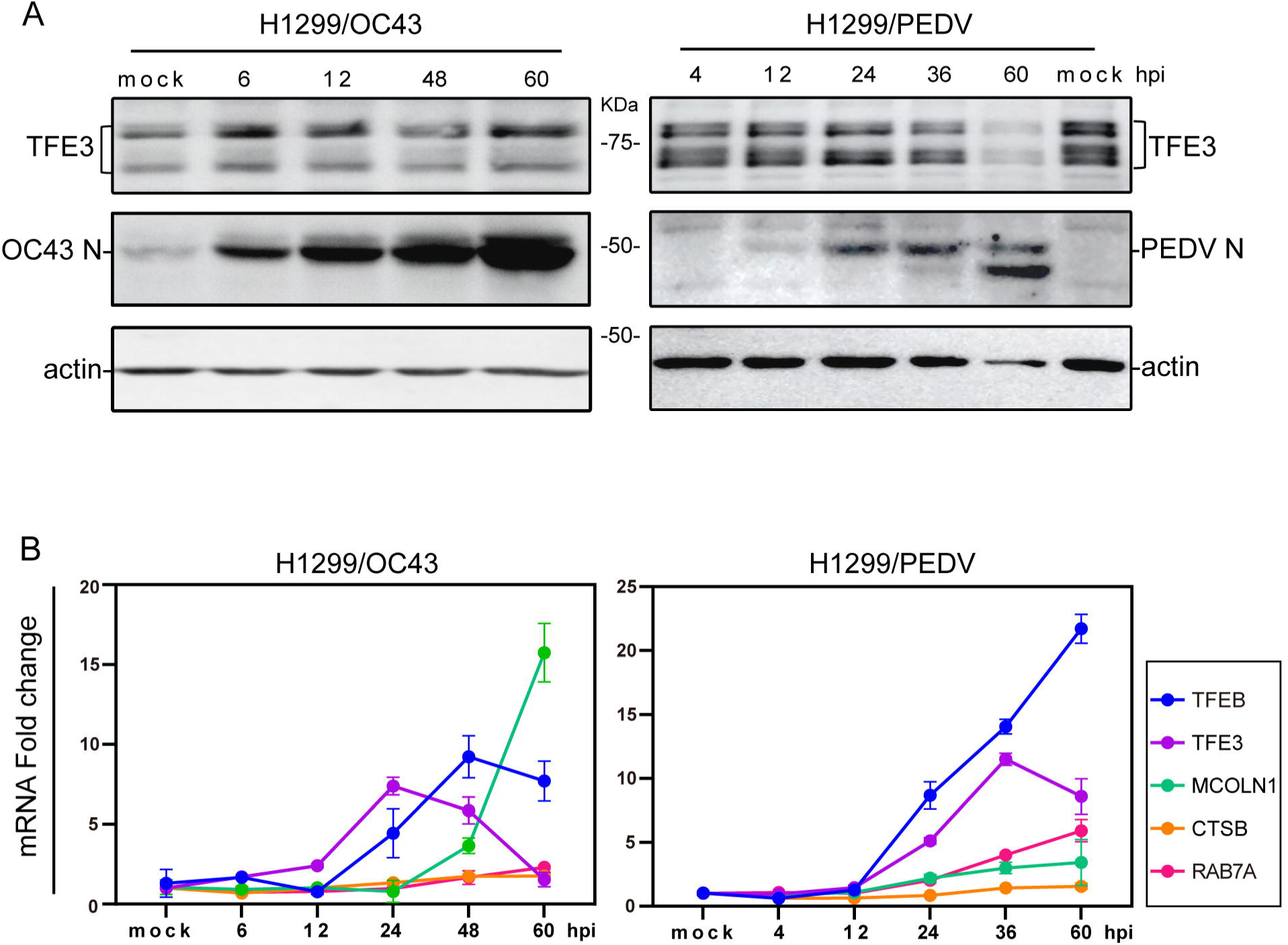
Induction of lysosomal stress response in culture cells infected with HCoV-OC43 and PEDV. A. H1299 cells were infected with HCoV-OC43 and PEDV, respectively, at an MOI∼2 or mock-treated with UV-inactivated viruses. Cells were harvested at the indicated time points and subjected to Western blot analysis using the indicated antibodies. Beta-actin was included as the loading control. B. Total RNA samples were extracted from cells in (A) and subjected to RT-qPCR. The levels of TFEB, TFE3, MCOLN1, CTSB and RAB7A were determined by the ΔΔCT method after normalization to the GAPDH mRNA level in cells treated with the corresponding UV-inactivated virus.

### Inhibition of IBV replication by TFEB/TFE3

To directly investigate if TFE3 and TFEB are functionally involved in the regulation of coronavirus replication, a H1299 cell clone with stable TFE3-knockdown (H1299-shTFE3 cells) was constructed, with a knockdown efficiency of about 70% (Fig. 4A-B), and a slightly but significantly reduced cell proliferation rate, as detected by CCK8 (Fig. 4C). Simultaneous knockdown of TFEB in this cell clone was also carried out with small interfering RNA (siRNA), achieving a knockdown efficiency of about 90% (Fig. 4E). Infection of the knockdown cells with IBV showed that knockdown of TFE3 facilitated IBV replication, and a similar trend effect was observed when both TFEB and TFE3 were simultaneously knocked down, as revealed by Western blot analysis of IBV N protein (Fig. 4D), and RT-qPCR analysis of viral genomic RNA in knockdown and control cells (Fig. 4E).

**Figure 4.**
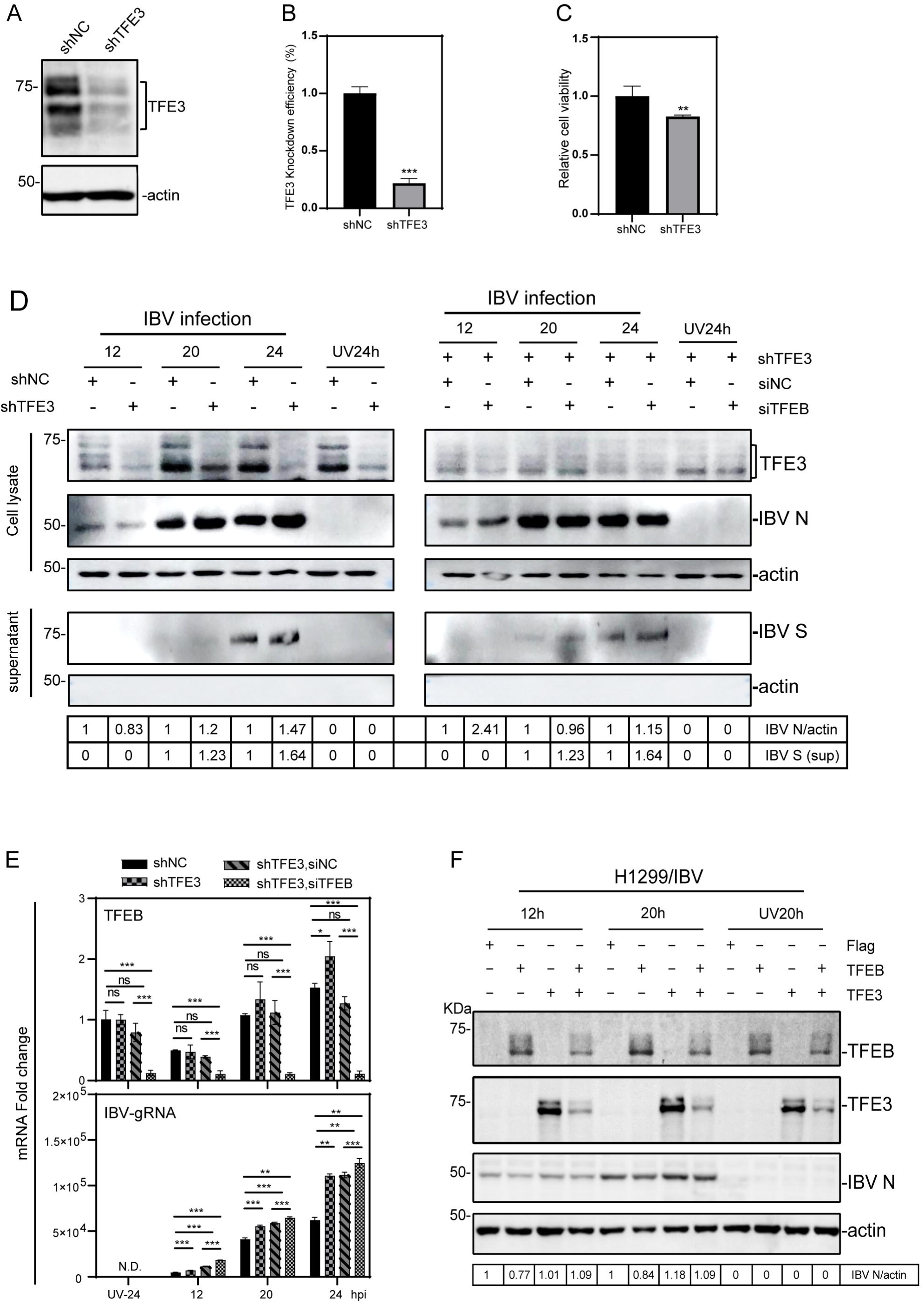
Regulation of viral replication by TFEB and TFE3 during IBV infection. A. Stable H1299-shTFE3 cell clone was established as described in Material and Method section. The knockdown efficiency was assessed by Western blot using antibodies specific for TFE3. Beta-actin was included as the loading control. B. The knockdown efficiency in stable H1299-shTFE3 cell clone described in (A) was assessed by RT-qPCR at the mRNA level. TFE3 mRNA level was determined by ΔΔCT method after normalization to GADPH in shNC cells (***, p < 0.001). C. The cell proliferation of stable TFE3-knockdown H1299 cell clones described in (A) was assessed by CCK8 (**, p < 0.01). D. The stable H1299-shTFE3 cells were either directly infected with IBV at an MOI∼2 or transfected with the indicated siRNA 24 hours prior to viral infection. Cells and cultured supernatants were harvested separately and subjected to Western blot using the indicated antibodies. E. The stable H1299-shTFE3 cells were transfected with the indicated siRNA, infected with IBV at an MOI∼2 and harvested at the indicated time points. Total RNA was extracted and mRNA levels of TFEB and IBV gRNA were determined by RT-qPCR (ns, non-significant; *, P < 0.05; **, p < 0.01; ***, p < 0.001). F. Effects of TFEB and/or TFE3 overexpression on the replication of IBV. H1299 cells were transfected with pXJ40-Flag, pXJ40-Flag-TFEB, pXJ40-Flag-TFE3, and pXJ40-Flag-TFEB+pXJ40-Flag-TFE3, respectively. At 24 hours post-transfection, cells were either infected with IBV at an MOI∼2 or treated with UV-inactivated IBV, harvested at the indicated time points and subjected to Western blot analysis using the indicated antibodies. Beta-actin was included as the loading control.

The effects of TFEB/TFE3 overexpression on IBV replication were then studied by cloning and overexpressing TFEB (mammalian) and TFE3 with an N-terminal Flag tag in H1299 (Fig. 4F) and TFEB (avian) in DF-1 cells (Supplementary Fig. S1), respectively. Compared with the empty vector, overexpression of TFE3 had little effect on IBV replication, but overexpression of TFEB assuredly reduced viral protein synthesis in both H1299 (Fig. 4F) and DF-1 cells (Supplementary Fig. S1). Coexpression of TFEB and TFE3 also reduced the expression of viral proteins, but at a less extent than that in cells overexpressing TFEB alone (Fig. 4F). In addition to the possibility that this Flag-tagged TFE3 may be functional inactive but act as a dominant negative regulator, the induction of the endogenous TFE3 expression at high levels in the infected cells may offset the functional effect of the overexpressed TFE3.

### Regulation of viral replication, autophagy and apoptosis by TFEB/TFE3 in cells infected with IBV, HCoV-OC43 and PEDV

The effect of TFEB/TFE3 knockdown on IBV-induced autophagy was then studied. SQSTM1, also known as p62, is a widely studied autophagy substrate, and its mRNA level can be used as a criterion for activating autophagy. The expression level of SQSTM1 in each group of IBV-infected cells was up-regulated with the increase of IBV infection time, compared with that in the UV-treated group. This is consistent with our previous report that IBV infection indeed induces autophagy (Fung and Liu, 2019b). In the knockdown cells infected with IBV, the induction level of SQSTM1 was significantly lower, compared with those in the control group, when TFE3 was knocked down alone or together with TFEB (Fig. 5A). These results demonstrate that upregulation of TFEB/TFE3 expression during IBV infection may be a strategy exploited by this virus to induce autophagy and regulate viral replication.

**Figure 5.**
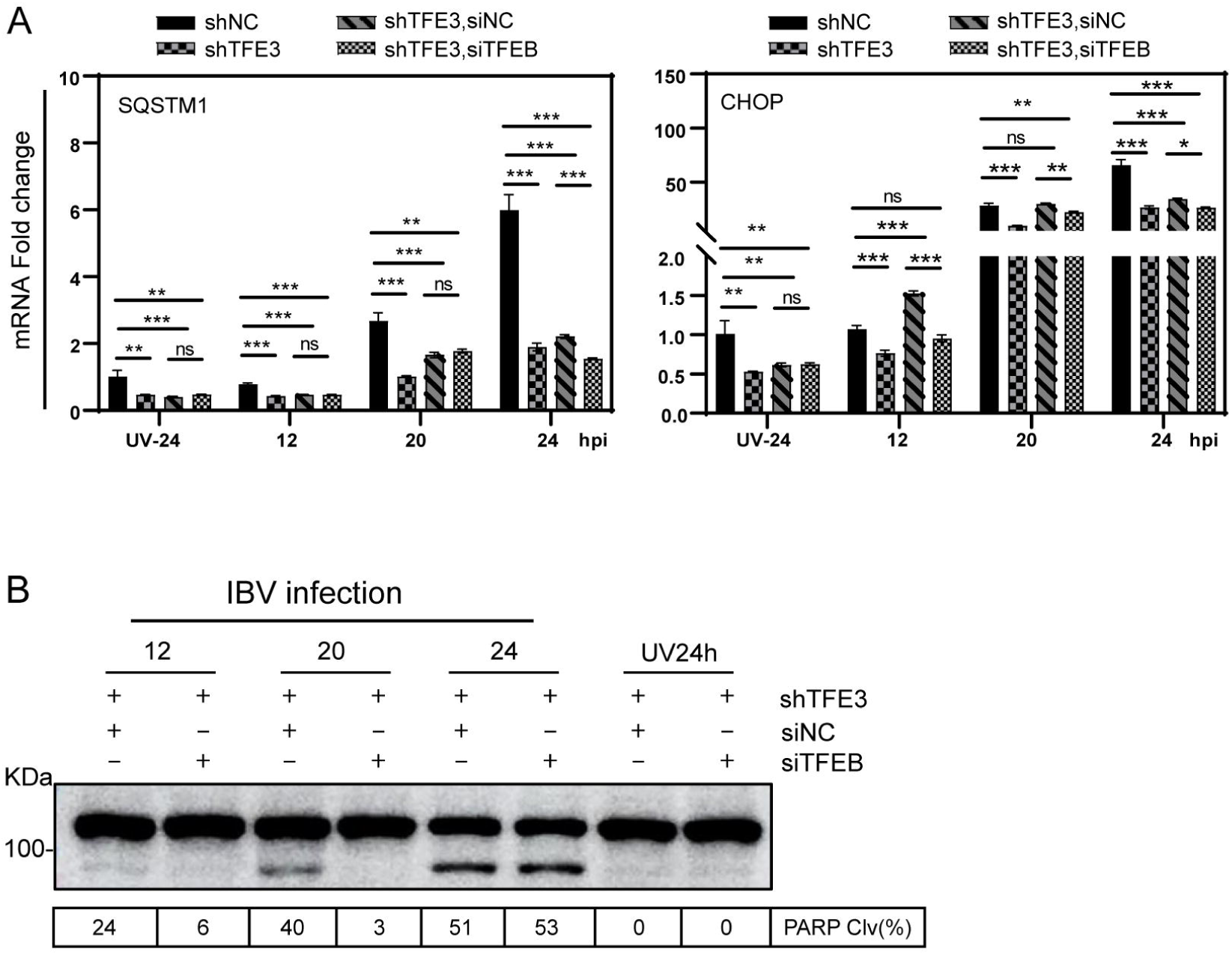
Regulation of autophagy and apoptosis by TFEB and TFE3 during IBV infection. A. Total RNA was extracted from IBV-infected stable H1299-shTFE3 cells described in Figure 4 and mRNA levels of SQSTM1 and CHOP were determined by RT-qPCR (ns, non-significant; *, P < 0.05; **, p < 0.01; ***, p < 0.001). B. Western blot analysis of the total cell lysates prepared from IBV-infected stable H1299-shTFE3 cells described in (A) using antibodies against PARP. The percentage of PARP cleavage [PARP Clv. (%)] was shown as the intensity of cleaved PARP (Cl) divided by the total intensities of the full-length PARP (FL) + Cl.

The effect of TFEB/TFE3 knockdown on IBV-induced apoptosis was assessed by determining the cleavage of poly(ADP-ribose) polymerase (PARP) as an apoptosis marker. Significantly lower percentage of PARP cleavage was detected in TFEB and TFE3 double-knockdown cells infected with IBV than that in the TFE3 single-knockdown cells at 12 and 20 hpi (Fig. 5B), suggesting the key role of these two factors in regulation of IBV-induced apoptosis.

CHOP, also known as GADD153, was previously shown to play an important regulatory role in endoplasmic reticulum (ER) stress-induced apoptosis during IBV infection (Liao et al., 2013). In a ChIP-seq analysis, CHOP and PUMA were identified as TFE3 targets, and both genes contained e-boxes in their promoter regions (Martina et al., 2016a). The inter-connection of these factors in regulation of IBV-induced apoptosis was then studied in TFE3 single-knockdown and TFEB/TFE3 double-knockdown cells. With increase of the infection time, the mRNA level of CHOP in each group of IBV-infected wild type cells was more significantly up-regulated, compared with that in cells treated with the UV-inactivated virus (Fig. 5A). This upregulateion was significantly inhibited by the knockdown of TFE3 alone or together with TFEB. As shown in Figure 5A, about 50% down-regulated CHOP expression was detected in the knockdown cells, compared with the negative control cells. These results demonstrate that IBV-induced TFEB/TFE3 upregulation may promote apoptosis by inducing ER stress response through upregulation of CHOP, thereby inhibiting viral replication.

Similar experiments in knockdown cells infected with HCoV-OC43 and PEDV, respectively, were performed to explore if TFEB/TFE3 may play similar regulatory roles in cells infected with these coronaviral and non-coronaviral viruses. In the knockdown cells infected with HCoV-OC43, the results were similar to those in IBV-infected cells. Knockdown of both genes promoted viral replication (Supplementary Fig. S2A-B) and inhibited the upregulation of SQSTM1, but higher levels of CHOP were detected in TFEB and TFE3 double-knockdown cells infected with HCoV-OC43 than those in the TFE3 single-knockdown cells at 12, 36 and 60 hpi (Supplementary Fig. S2B). In cells infected with PEDV, knockdown of TFE3 promoted viral replication (Supplementary Fig. S3A-B), but very limited additional effects on viral replication, autophagy and apoptosis were observed in the double-knockdown cells, compared with knockdown of TFE3 alone (Supplementary Fig. S3B). Taken together, these results demonstrate that TFEB/TFE3 may play general roles in regulation of viral replication and virus-induced apoptosis and autophagy.

### Implication of the lysosomal pathway in the release of HCoV-OC43 virions

Recent studies have demonstrated that betacoronavirus MHV excretes cells using the lysosomal transport, unlike other enveloped RNA viruses either using the secretion pathway or through fusion with the plasma membrane and budding (Ghosh et al., 2020). To investigate if the lysosomal pathway is a releasing mechanism utilized by IBV, HCoV-OC43 and PEDV, Brefeldin A (BFA) was added to inhibit the biosynthetic secretory pathway. The effects of different concentrations of BFA on cell viability were first checked to rule out the possible detrimental effects of high concentrations of BFA on the cell proliferation. Different concentration gradients of BFA were added to H1299-shTFE3 cells, and the cell viability was measured with CCK8, showing that the cell viability was not affected by adding 0-10 μg/mL of BFA (Fig. 6A). Virus-infected cells were then treated with 5 μg/mL of BFA, replaced with fresh medium at 10 hours post-BFA addition, and both cells and supernatants were collected at indicated time points for Western blot analysis. Compared with cells incubated with the same amount of DMSO at the same time point, in cells infected with IBV and PEDV, viral proteins were readily detected in cell lysates after the addition of BFA, but IBV S and PEDV S were completely undetectable in the supernatants, indicating that the addition of BFA completely blocked the release of IBV and PEDV particles (Fig. 6B&C). However, viral proteins were detected in the supernatants in cells infected with HCoV-OC43, after treatment of cells with BFA, (Fig. 6D). These results demonstrated that the lysosomal pathway may be involved in the release of HCoV-OC43, but not IBV and PEDV particles.

**Figure 6.**
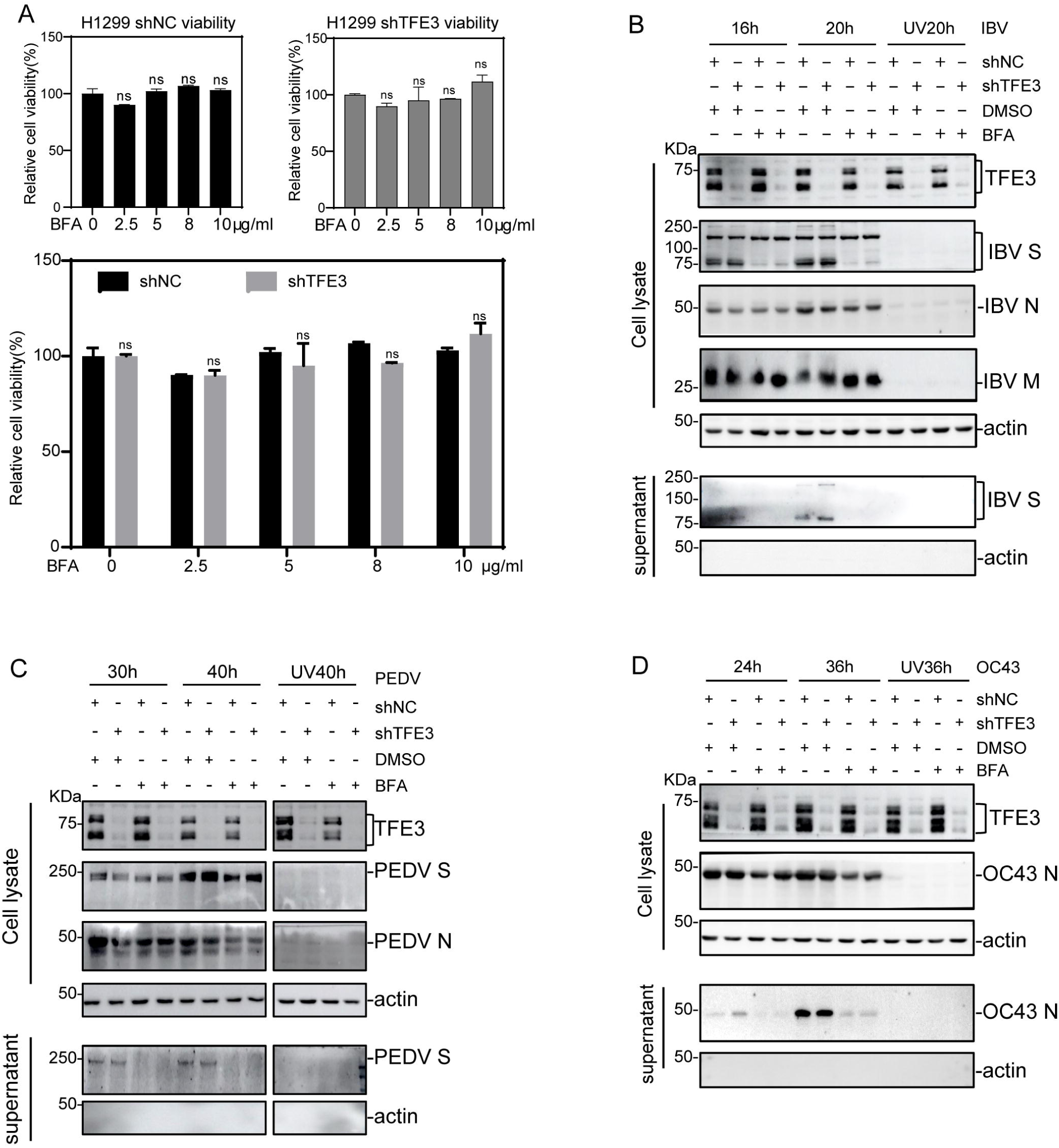
Differential roles of the lysosomal pathway in the release of coronavirus particles. A. Effects of different concentrations of BFA on the proliferation of the stable TFE3-knockdown H1299 cell clones. Cells were cultured in 96-well plates overnight, medium. At 24 hours post-BFA treatment, CCK8 was added and the absorbance at 450nm was measured after incubation for 2 hours. The cell proliferation rates were calculated after normalization to the absorbance value in cells incubated with DMSO only (ns, non-significant; *, P < 0.05; **, p < 0.01; ***, p < 0.001). B/C/D. The stable TFE3-knockdown H1299 cells were infected with IBV, PEDV and HCoV-OC43, respectively, at an MOI∼2, and 5μg/ml BFA were added 10 hours post-infection. Total cells and supernatants were harvested separately at the indicated time points and were analyzed by Western blot with the indicated antibodies.

## Discussion

Viral replication cycle is closely associated with host cell factors and signaling pathways, imposing drastic impacts on the cellular structure and physiology of different organelles. This would lead to the activation of host cell stress responses, which in turn regulate viral replication, pathogenesis, virus-induced apoptosis, autophagy and innate immunity. Lysosomes are now considered key players in regulation of these cellular processes. In this study, we report the upregulation of TFEB/TFE3 as well as downstream targets and induction of lysosomal stress response in cells and/or chickens infected with three coronaviruses of different genera and a non-coronavirus. TFE3 was shown to play a prominent role in regulating viral replication, virus-induced autophagy and apoptosis in these infected cells, and a synergistic effect with TFEB in IBV- and HCoV-OC43-infected cells. Inhibition of the biosynthetic secretory pathway with BFA revealed lysosomal pathway-dependent release of mature HCoV-OC43 particles. Figure 7 summarizes the key lysosomal stress-related signaling pathways and their regulatory roles in the replication of these viruses.

**Figure 7.**
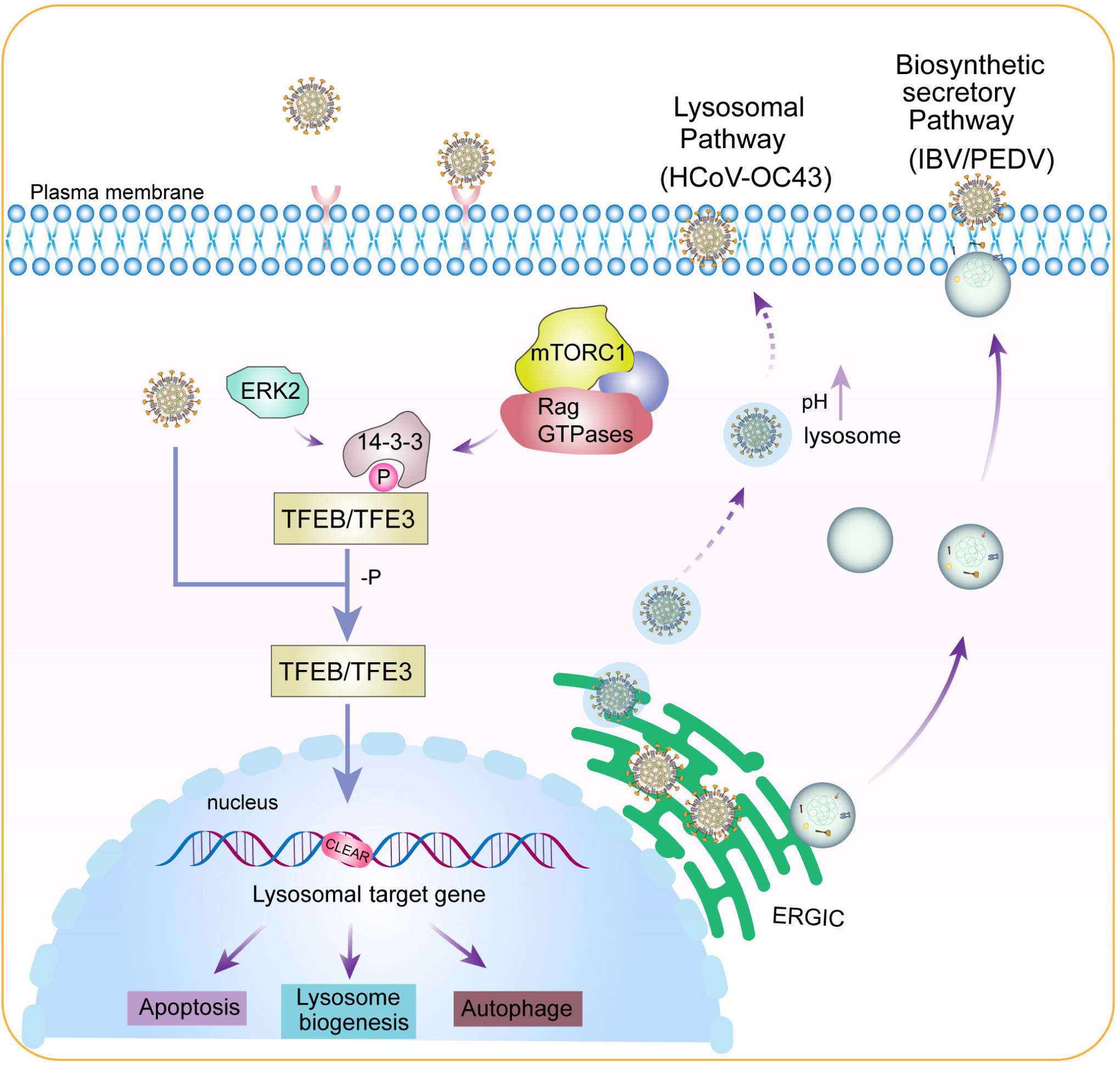
Diagram illustrating the current working model. Pointed arrows denote activation, P and -P denote phosphorylation and dephosphorylation, respectively, and dotted lines denote processes that are not fully characterized.

Autophagy is a self-protection mechanism that is closely related to apoptosis. It normally occurs before, but independently of, apoptosis (Yu et al., 2018). This sequential occurrence of autophagy and apoptosis may serve two purposes. First, autophagy may be induced by dying cells to initiate their catabolism, thus accelerating the clearance of dying cells. Second, autophagy may also help maintain optimally high ATP levels, which may lead to apoptosis when cells are unable to achieve homeostasis and are subjected to constant stress (Maiuri et al., 2007). TFE3 was shown to be a powerful regulator controlling the expression of autophagy flux-related genes in a variety of cancers. Studies have shown that TFE3 can enhance the number and function of lysosomes, enhanced autophagy, and promote the pathological process of pancreatic ductal adenocarcinoma (Perera et al., 2015). Martina et al. found that TFEB- and TFE3-knockout MEF cells were significantly less susceptible to ER stress-induced cell death (Martina et al., 2016a). ER dysfunction was caused by transcriptional up-regulation of pro-apoptotic factors such as CHOP and PUMA(direct targets of CHOP and ATF4) to promote cell apoptosis. TFEB and TFE3 may, therefore, regulate the expression of CHOP and PUMA by inducing ATF4 expression or directly binding to their promoters, promoting apoptosis under prolonged ER stress conditions by maintaining sustained ATF4 activation (Martina et al., 2016b). In this study, infection of TFEB- and TFE3-knockdown cells with IBV, HCoV-OC43 and PEDV showed down-regulation of the pro-apoptotic factor CHOP and autophagy-related gene SQSTM1, revealing the enhancement roles of TFEB and TFE3 in coronavirus-induced autophagy and apoptosis.

Elucidation of the regulatory roles of lysosomes and lysosomal biogenesis in innate immunity against viral infection is beginning to emerge. During dendritic cell maturation, TFEB activation enhances phagosome acidification, increases protein degradation, and improves major histocompatibility complex (MHC) class II antigen presentation, all of which are critical for initiating T cell responses against viral infections(Samie and Cresswell, 2015). Shortly after macrophages are infected with HIV, TFEB is activated via a TLR8-dependent pathway, leading to a brief increase in autophagy and enhancement of HIV replication (Campbell and Spector, 2013). Our results presented here also support that activation of lysosomal biogenesis in cells infected with three different coronaviruses and a non-coronavirus accelerates clearance of these virus after infection.

As final steps in the viral replication cycle, assembly and release of mature virion particles are highly complex and dynamic processes. It is generally believed that the assembly and germination of coronaviruses occur in the ERGIC, soon after the newly synthesized viral genomic RNA coated with N protein enters the ERGIC through budding. The virus particles are then enveloped by host membranes containing viral M, E, and S transmembrane structural proteins. Once in ER/ERGIC, the virus particles enter the Golgi apparatus and the trans-Golgi network (TGN) for maturation by glycosylation and other post-translational modifications, transported in smooth vesicles through the biosynthetic secretory pathways, and released by binding of the vesicles to the cytoplasmic membrane to expel the viral particles from the cell (Fung and Liu, 2019a). This viral shedding process is shared by other enveloped RNA viruses, such as HCV, dengue virus and West Nile virus (Ravindran et al., 2016; Robinson et al., 2018), and was thought to be the only pathway used by all coronaviruses for virion release. By using BFA to block the secretory pathways, recent studies, however, have provided evidence that betacoronavirus MHV uses the lysosomal transport to excrete cells. The blockade of the secretory pathway by BFA may be manifested in two ways, one at the level of the ER-Golgi and the other at the trans-Golgi network. However, transport from the Golgi complex and the plasma membrane to the lysosome remains unaffected. (Strous et al., 1993). Our observations that only the mature HCoV-OC43 particles were efficiently released in the presence of 5 μg/mL of BFA demonstrated that these two viruses may be released through transport from lysosome to plasma membrane in addition to secretory pathways, while IBV and PEDV may only use the secretory pathways. Why do viruses use lysosomes to escape cells without being digested? One study using specific inhibitors to inhibit lysosomal acidification found that the release of infectious HCV particles was greatly reduced (Wozniak et al., 2010). Deacidification of lysosomes in infected cells, a general phenomenon observed in cells infected with IBV, PEDV, HCoV-OC43 (unpublished observations), would reduce the protease activities and facilitate the release of these viruses.

In conclusion, this study provides novel insights into the functional roles of the lysosomal biogenesis and induction of lysosomal stress response during infection of cells and/or chickens by three coronaviruses of different genera and a non-coronavirus. This would pave a way for further studies of the underlying mechanisms and signaling pathways in regulation of coronavirus replication and virus-host interactions, identifying novel lysosome-related cellular targets for developing antiviral intervention.

## Acknowledgement

This work was partially supported by the National Natural Science Foundation of China (grant number 32170152), Guangdong Basic and Applied Basic Research Foundation of China (grant number 2024A1515012930), and Zhaoqing Xijiang Innovative Team Foundation of China (grant numbers P20211154-0201 and P20211154-0202).

