## Supplementary figures and images for "Modulation of viral replication, autophagy and apoptosis by induction and mutual regulation of transcription factors EB and E3 during coronavirus infection"

### Supplemental figure 1

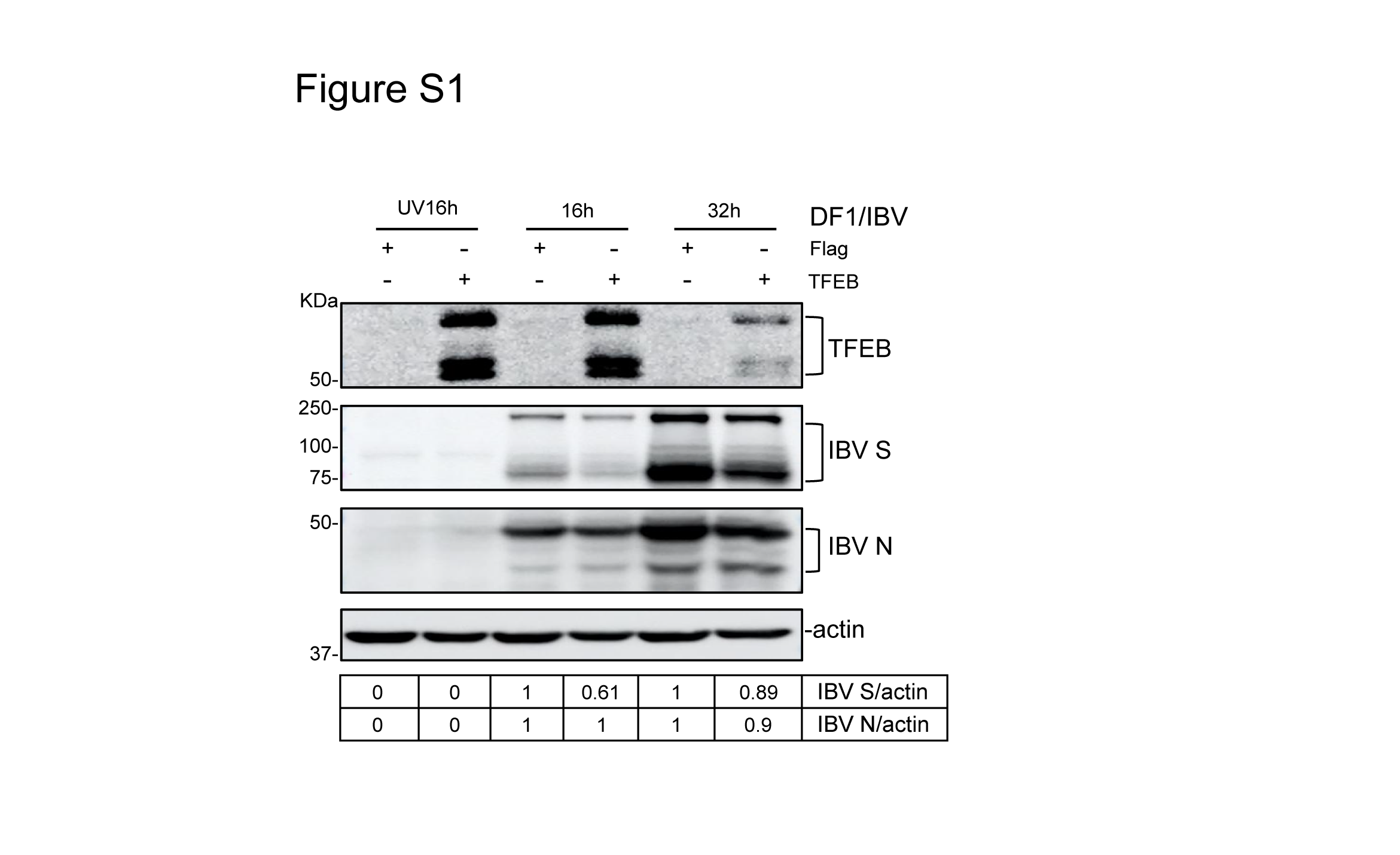

### Supplemental figure 2

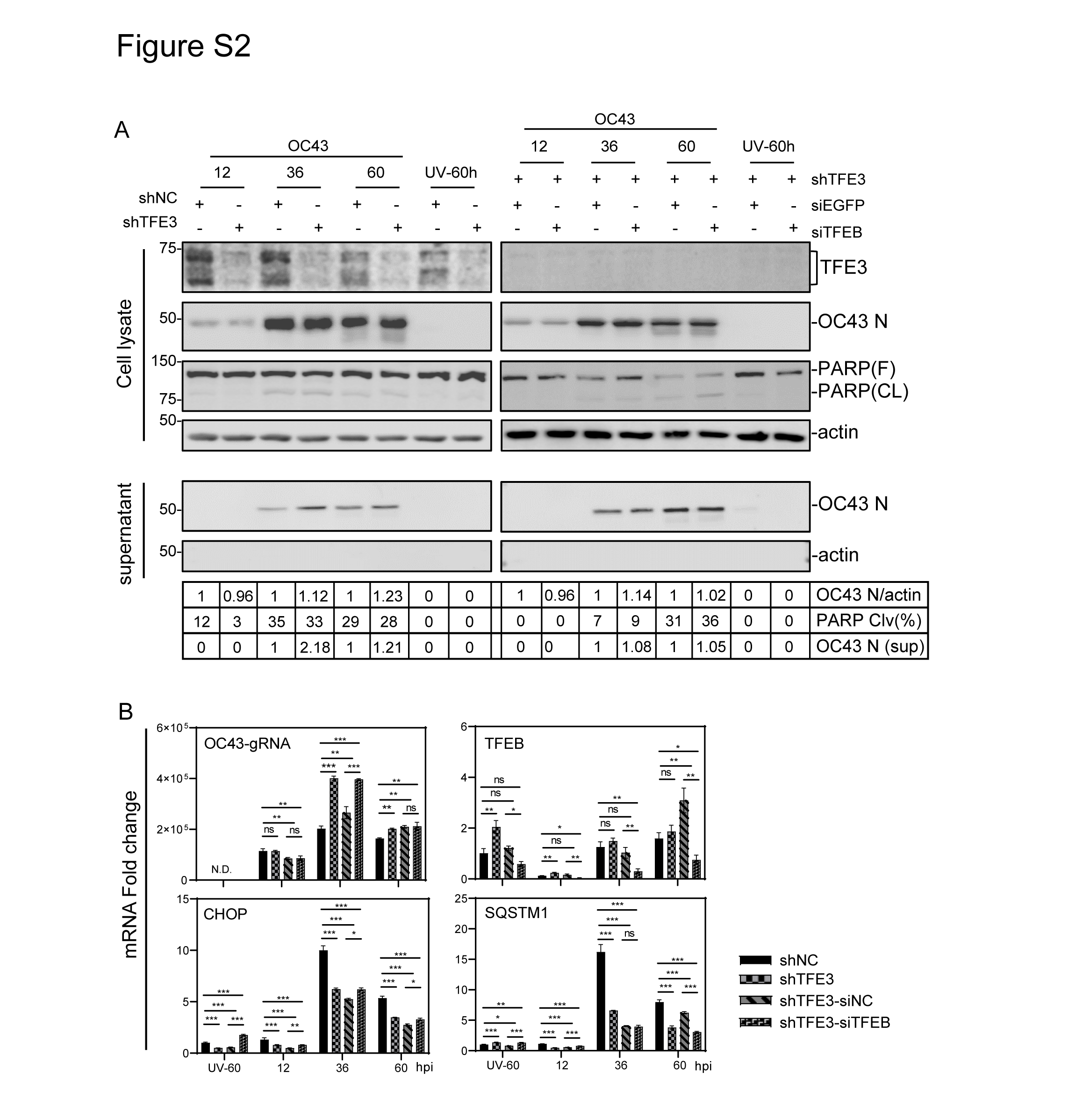

### Supplemental figure 3

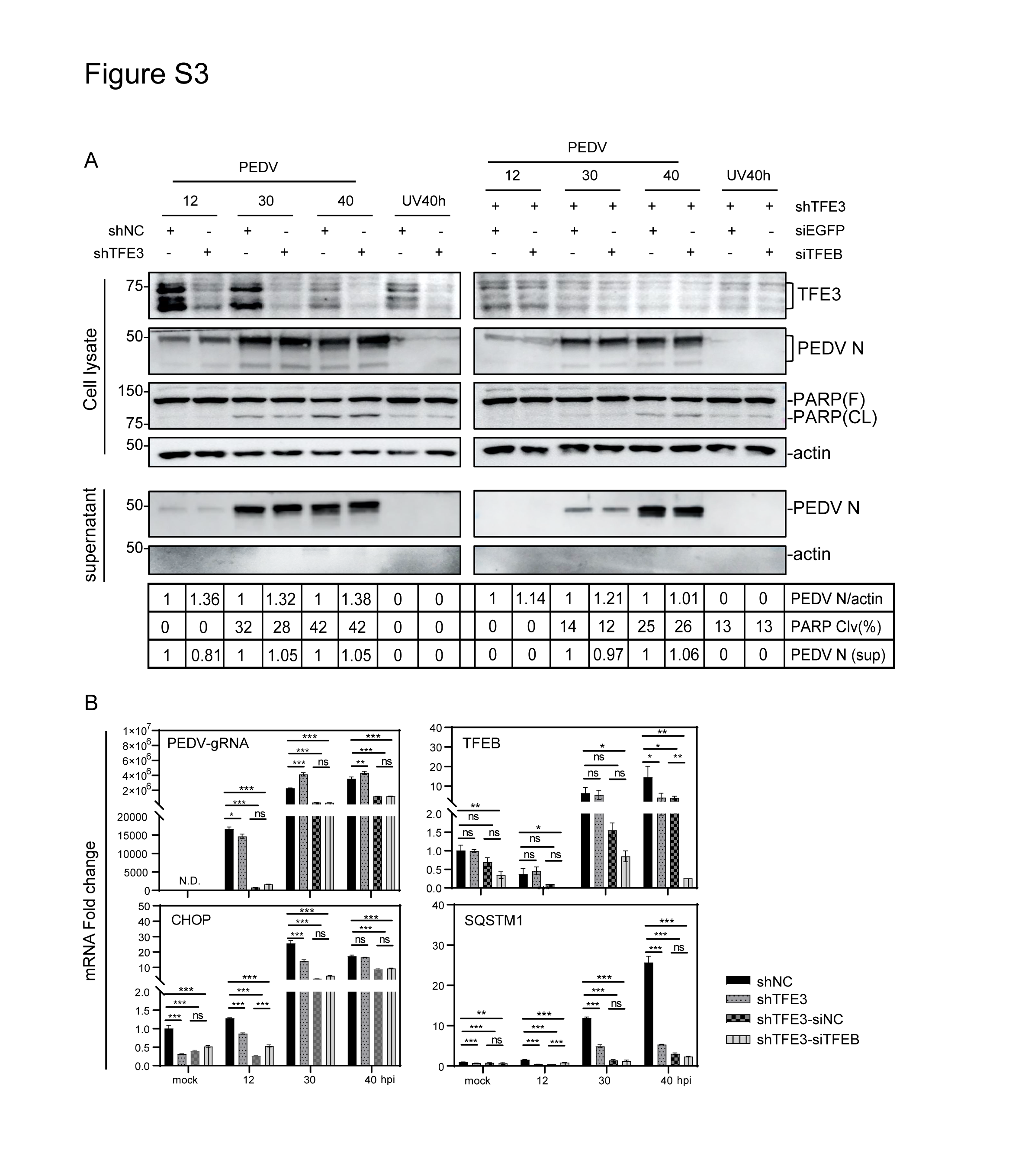
